# The CCCTC-binding factor CTCF represses hepatitis B virus Enhancer I and regulates viral transcription

**DOI:** 10.1101/2020.05.08.085548

**Authors:** V D’Arienzo, J Ferguson, G Giraud, F Chapus, JM Harris, PAC Wing, A Claydon, S Begum, X Zhuang, P Balfe, B Testoni, JA McKeating, JL Parish

## Abstract

Hepatitis B virus (HBV) infection is of global importance with over 2 billion people exposed to the virus during their lifetime and at risk of progressive liver disease, cirrhosis and hepatocellular carcinoma. HBV is a member of the *hepadnaviridae* family that replicates via episomal copies of a covalently closed circular DNA (cccDNA) genome. The chromatinization of this small viral genome, with overlapping open reading frames and regulatory elements, suggests an important role for epigenetic pathways to regulate viral transcription. The chromatin-organising transcriptional insulator protein CCCTC-binding factor (CTCF) has been reported to regulate transcription in a diverse range of viruses. We identified two conserved CTCF binding sites in the HBV genome within Enhancer I and chromatin immunoprecipitation (ChIP) analysis demonstrated an enrichment of CTCF binding to integrated or episomal copies of the viral genome. siRNA knockdown of CTCF results in a significant increase in pre-genomic RNA levels in *de novo* infected HepG2 cells and those supporting episomal HBV DNA replication. Furthermore, mutation of these sites in HBV DNA minicircles abrogated CTCF binding and increased pre-genomic RNA levels, providing evidence of a direct role for CTCF in repressing HBV transcription.

**IMPORTANCE:** Hepatitis B virus (HBV) is a global cause of liver disease. At least 300 million individuals are chronically infected with HBV, frequently leading to life-threatening liver cirrhosis and cancer. Following viral entry, HBV DNA enters the nucleus and is bound by histones that are subject to epigenetic modification. The HBV genome contains two enhancer elements that stimulate viral transcription but the interplay between the viral enhancers and promoters is not fully understood. We have identified the host cell protein CCCTC binding factor (CTCF) as a repressor of HBV gene expression. CTCF binds to the HBV genome within Enhancer I and represses transcription of pre-genomic RNA. These findings provide new insights into how HBV transcription is regulated and show a new role for CTCF as a transcriptional insulator by associating with the viral genome between Enhancer I and the downstream basal core promoter.

## INTRODUCTION

Hepatitis B virus (HBV) infection is one of the world’s unconquered infections with an estimated 2 billion people exposed to the virus in their lifetime. HBV replicates in hepatocytes and chronic infection can result in progressive liver disease, cirrhosis and hepatocellular carcinoma (HCC). HBV is a member of the *hepadnaviridae* family and classified into eight genotypes, A-H, based on a sequence divergence of greater than 8% (1, 2). Viral genotypes are associated with differences in clinical outcome and treatment responses (3, 4). The HBV genome is a small, partially double-stranded relaxed circular DNA (rcDNA) genome of approximately 3.2 Kb. Following HBV entry into hepatocytes via the liver-specific bile-acid transporter, sodium taurocholate co-transporting polypeptide (NTCP) (5, 6), rcDNA is released into the nucleus and is repaired into covalently closed circular DNA (cccDNA). cccDNA persists in the nucleus as multiple copies of nucleosome-associated minichromosomes which serve as a template for virus transcription (7). Establishment of the stable long-lived cccDNA intermediate is thought to be responsible for persistence of HBV infection (8, 9).

The HBV genome is transcribed by the host RNA polymerase II (RNA pol II) complex from unique promoters (basal core promoter (BCP), Sp1, Sp2 and Xp) and transcription start sites (7). This results in the generation of six major viral RNAs of increasing length with heterogeneous 5’ ends and a common polyadenylation signal (10, 11). The 3.5kb preC transcript encodes pre-core or e antigen (HBe) protein. Approximately 100 base pairs downstream is the transcriptional start site for pre-genomic (pg) RNA which encodes the core (HBc) protein and the viral polymerase (pol). When encapsidated in the cytoplasm, pgRNA forms the template for the reverse-transcription of new rcDNA molecules by the viral pol (7). The large, medium and small surface envelope proteins (HBs) are encoded by the 2.4 kb preS1 and 2.1 kb preS2/S transcripts. The smallest transcript is the 0.7kb X RNA which encodes the hepatitis B virus x protein (HBx) protein, which has been shown to influence many host cell pathways including regulation of transcription of viral and host genes, metabolism and cell cycle. Two viral enhancers play an important role in the regulation of HBV transcription. Enhancer I (EnhI) is located upstream of and partially overlaps the X promoter (Xp) and regulates transcription of HBx and core genes. It also directs basal core promoter (BCP) activity (12), which stimulates the production of both preC and pgRNAs (13). Enhancer II (EnhII) overlaps a large portion of BCP and functions to stimulate activity of the distal Sp1 and Sp2 promoters as well as Xp and BCP (14). The BCP encodes a negative regulatory element (NRE) that overlaps with Enh II (15) and has been reported to repress EnhII-mediated promoter activation (16).

Nuclear HBV cccDNA is assembled into nucleosomes by cellular histones to form episomal chromatin (17, 18). The viral DNA is enriched with active epigenetic histone modifications including trimethylation of lysine 4 (H3K4Me3) and acetylation of lysine 27 on histone 3 (H3K27Ac) but devoid of the repressive marks such as trimethylation of lysine 27 on histone 3 (H3K27Me3) (19, 20). The overlap of active histone marks with RNA pol II occupancy suggests that viral transcription is regulated by epigenetic modification. In support of this, treating *de novo* infected primary human hepatocytes with inhibitors of the histone acetyltransferase p300/CBP reduces HBV RNA levels (19). Although the mechanisms underlying the epigenetic regulation of HBV cccDNA are not fully understood (21), several epigenetic modifiers are recruited to HBV cccDNA by HBx. As such, HBx behaves as a transcriptional regulator of both viral and cellular promoters (22) and although HBx cannot bind to DNA directly, it can associate with components of the basal transcription machinery, transcription factors and transcriptional co-activators (23). HBx coordinates the recruitment of the CBP/p300 and PCAF histone acetyl transferases (HAT) to cccDNA while facilitating the exclusion of histone deacetylases (HDACs) HDAC1 and Sirtuin 1 (Sirt1), resulting in hyperacetylation of cccDNA (24, 25). HBV transcription is dependent on an array of ubiquitous and liver-specific cellular transcription factors including the liver specific hepatocyte nuclear factors 1 and 4 (HNF-1/4) and ubiquitously expressed octamer-binding protein 1 (Oct-1) and specificity protein 1 (SP1) (26).

The genomes of metazoans are highly organised into megabase-sized regions termed topologically-associated domains (TADs) that provide regulatory segmentation required for appropriate gene expression and replication. TADs are separated by regions enriched in binding sites of the ubiquitously expressed CCCTC binding factor (CTCF) which stabilises chromatin loops by anchoring cohesin rings at the base of the loops (27). Such spatial organisation can create epigenetic boundaries that separate transcriptionally active and inactive chromatin domains and control cis-regulatory elements such as transcriptional enhancers. CTCF binds to tens of thousands of either ubiquitous or cell type specific consensus binding sites within the human genome, regulating both tissue-specific and developmental changes in gene expression (28).

The occupancy of specific CTCF binding sites is dictated by chromatin accessibility and local epigenetic status (29). In addition to the organisation of chromatin domains, CTCF can function as a transcriptional repressor or activator by direct association with promoter proximal elements. CTCF was shown to act as a transcriptional repressor of the *c-myc* oncogene by creating a roadblock to RNA pol II (30). Conversely, CTCF can physically associate with transcriptional regulators such as the general transcription factor, TFII-I to promote recruitment of the cyclin dependent kinase 8 (CDK8) resulting in stimulation of RNA pol II activity (31). CTCF regulates the transcription (up or down) of evolutionarily distinct DNA viruses (32) including: Kaposi sarcoma-associated herpesvirus; Epstein-Barr virus and herpes simplex virus (33-38). We have demonstrated that CTCF recruitment to the human papillomavirus (HPV) genome negatively regulates early promoter usage via host cell differentiation-specific stabilisation of an epigenetically repressed chromatin loop (39, 40). However, a role in HBV transcription regulation has not yet been reported, herein we show that CTCF binds HBV DNA and acts as a repressor of viral transcription.

## RESULTS

### CTCF binds HBV DNA at conserved sites within enhancer elements

To assess whether CTCF binds HBV DNA, we selected two independent ‘HBV producer’ HepG2 lines, HepG2.2.15 (41) and HepAD38 (42) that carry integrated copies of 1.3x overlength HBV genomes, maintain cccDNA and generate infectious virus. We isolated and sheared chromatin from nuclear fractions to limit contamination of cytoplasmic rcDNA and performed an anti-CTCF chromatin immunoprecipitation (ChIP) followed by quantitative PCR (ChIP-qPCR). Primers were selected to amplify 100-200 base pair regions of the HBV genome to provisionally identify CTCF binding sites. We show low level CTCF binding above the control IgG across the viral DNA with a significant enrichment in the Xp region in both cell lines **(Fig.1A)**. Analysing histone modifications of HBV chromatin from HepG2.2.15 cells showed the viral DNA lacked the repressive H3K27Me3, in agreement with previous reports (19, 20) **(Fig.1B)**. ChIP for histone marks associating with active transcription, including H4Ac and H3K4Me3, identified these epigenetic marks throughout the viral genome, with an enrichment in the BCP and Xp regions. Since both HepG2.2.15 and HepAD38 cell lines have cccDNA and integrated viral genomes, we are unable to discriminate CTCF binding between these forms of viral DNA. We therefore studied HepG2 cells expressing an episomal copy of HBV DNA (HepG2-HBV-Epi) (43) to establish whether CTCF can bind episomal viral DNA. We first assessed whether episomal copies of HBV DNA are sheared by sonication by PCR amplification of viral targets of increasing length pre- and post-sonication. While the unsheared chromatin yielded a series of PCR products of increasing length, only amplicons below 238 base pairs were detected in the sonicated material **(Fig.1C)**. Amplicons over 353 base pairs were barely visible in the sonicated samples, demonstrating effective shearing of episomal HBV genomes. ChIP of sheared chromatin isolated from HepG2-HBV-Epi nuclear extracts showed CTCF bound to the EnhI region of the viral DNA **(Fig.1D).** We noted relatively lower ChIP of viral DNA from the HepG2-HBV-Epi cells compared to HepG2.2.15 or HepAD38 cells, this may reflect differences in the epigenetic status of the viral DNA in these model systems. Our observation that CTCF binds to EnhI, the major transcriptional regulatory element of the BCP and Xp, suggests that CTCF regulates its activity.

**Figure 1:**
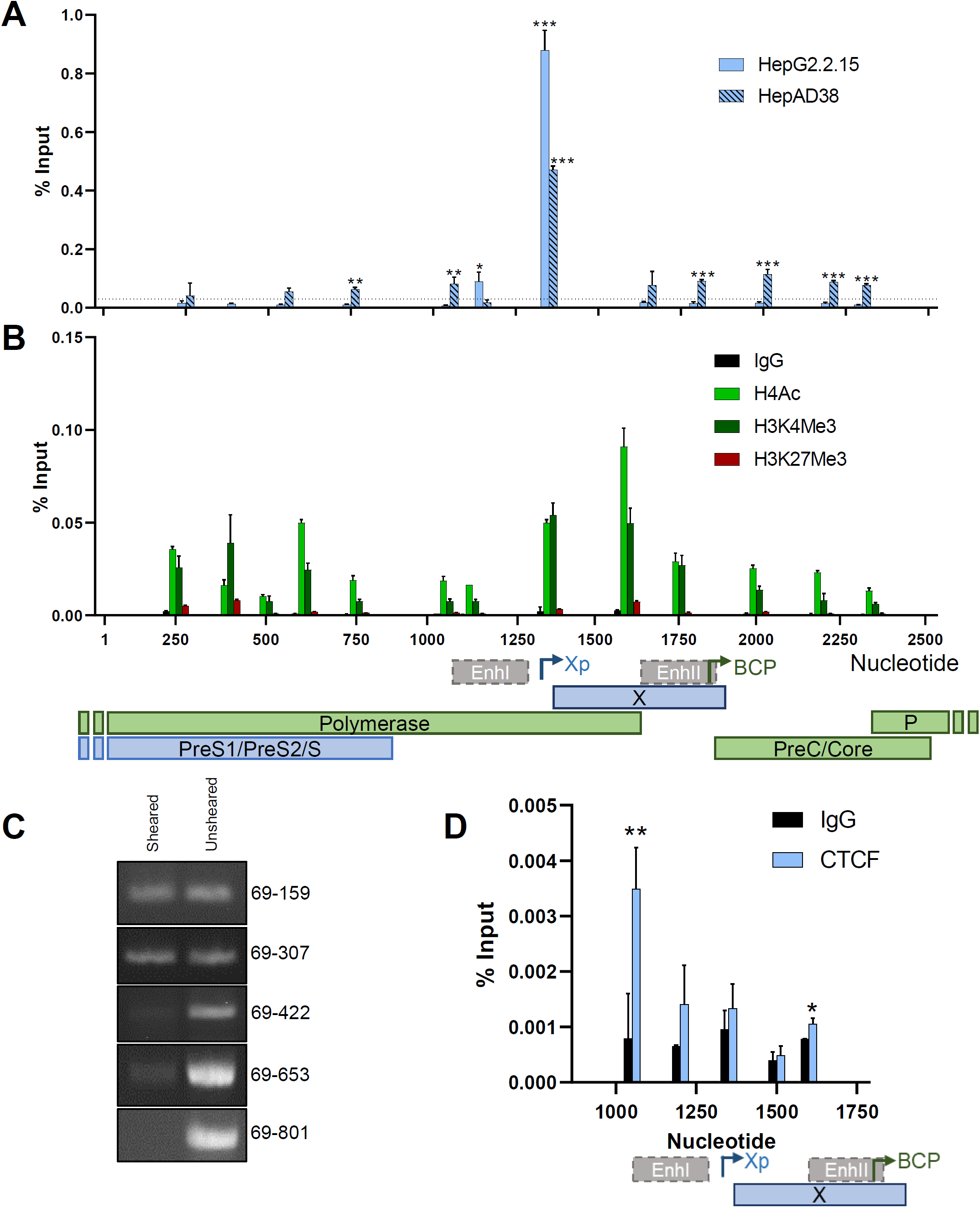
CTCF associates with HBV DNA and is enriched at viral Enhancer I and X promoter. (A) Association of CTCF with HBV DNA in HepG2.2.15 and HepAD38 cells was analysed by ChIP-qPCR and presented as % Input recovery. Statistical significance shows comparison of CTCF-specific ChIP with maximal recovery using IgG control (dotted line). (B) The distribution of histone modifications (H3K4Me3, H3K27Me3 and H4Ac) in HepG2.2.15 cells by ChIP-qPCR. (C) Efficiency of chromatin shearing in HepG2-HBV-Epi cells was assessed by PCR of sonicated versus non-sonicated chromatin. Amplicons were generated with a constant sense primer (anneals at nt 69) and anti-sense primers binding at increasing distance from the sense primer (nt 159, 307, 422, 653 and 801). Amplification of HBV DNA was assessed by SyBr green staining of bands separated by electrophoresis. (D) Association of CTCF was assessed by ChIP-qPCR. (A, B and D) Data shown are the mean +/- SEM of three technical repeats and are representative of three biological repetitions. P values were determined using a paired t test. *denotes p <0.05, **denotes p <0.01, ***denotes p <0.001. Annotation of HBV genome features including open reading frames, enhancers and selected promoters is shown below the histograms.

HepG2.2.15, HepAD38 and HepG2-HBV-Epi cells contain HBV genotype D and having demonstrated that CTCF associates with viral DNA in all three cell lines, we used an open access CTCF binding site database (http://insulatordb.uthsc.edu/) to identify putative CTCF binding sites within HBV genotype D. We identified two CTCF binding sites (BS) between nucleotides 1194-1209 in EnhI (CTCF BS1) and 1275-1291 in the Xp (CTCF BS2), consistent with the single binding peak observed in our ChIP-qPCR analysis. Importantly, these binding sites are conserved amongst all HBV genotypes (>7000 sequences in HBV database [HBVdb.fr]) **(Fig.2A)**. The *hepadnaviridae* family includes a number of related viruses that infect other species including birds, mammals, fish, reptiles and amphibia. Inspection of reference sequences from distinct *hepadnaviridae* showed that both consensus CTCF binding sites are conserved in viruses infecting primates and the majority of mammals and bats but are absent from viruses infecting birds, fish or amphibia, demonstrating evolutionary conservation of both CTCF binding sites **(Fig.2B).**

**Figure 2:**
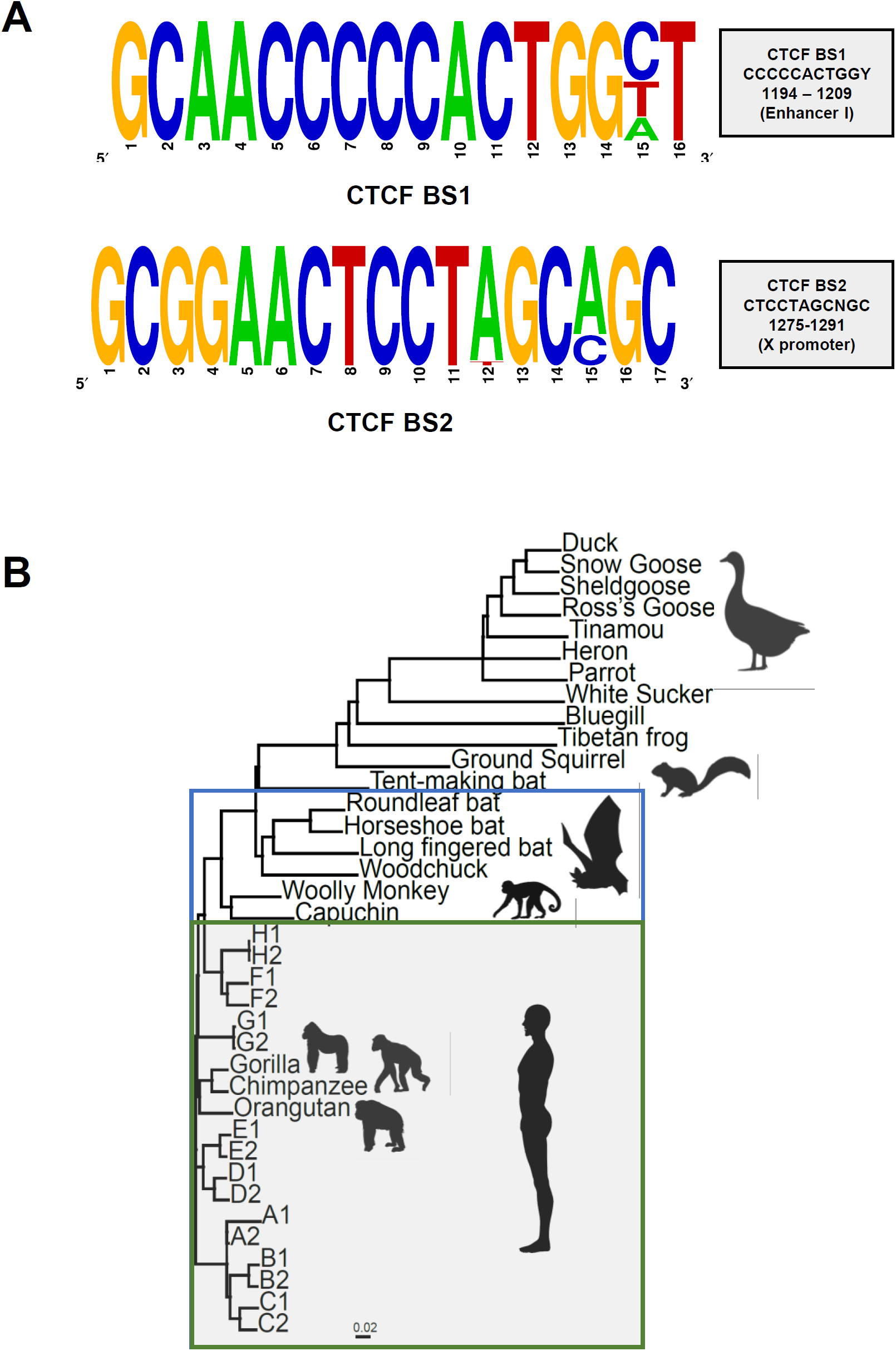
Identification of conserved CTCF binding sites in HBV genomes and diverse *hepadnaviridae*. (A) Conservation of CTCF BS among 7,313 HBV sequences (HBVdb.fr). All sites, except where indicated, are > 98% conserved. (B) Neighbor-joining phylogenetic tree of members of the *hepadnaviridae* (adapted from (1)). The green box shows viral genomes that encode both CTCF BS1 and 2 (all human and old world primate viruses), whereas the blue box shows viral genomes encoding only CTCF BS1 (new world monkeys, woodchucks and all bats except the tent making bat).

### CTCF represses HBV Enhancer I

To analyse the role of CTCF in regulating HBV enhancer activity, we used promoter constructs that encode Firefly luciferase under the control of the BCP (nt 900-1859) or EnhI and Xp (nt 900-1358) **(Fig.3A)** (44). We noted that the BCP showed a significantly lower (4-fold) activity compared to the EnhI construct, most likely reflecting the presence of a NRE at nt1613-1636 that can repress BCP activity **(Fig.3B)**. To analyse the function of CTCF in regulating EnhI activity, we silenced CTCF in HepG2-NTCP using an siRNA Smartpool **(Fig.3C)**, transfected the viral promoter plasmids along with a *Renilla* luciferase control plasmid and measured activity after 72h. Knockdown of CTCF protein increased EnhI activity, however we noted a minimal effect on the BCP activity, suggesting that CTCF represses EnhI but this effect is limited in the presence of an NRE in the full transcriptional reporter construct **(Fig.3D)**. To assess whether the putative CTCF BS mediated the control of EnhI, we introduced silent mutations into the pEnhI-Luc designed to abrogate CTCF binding (45), without altering the polymerase protein sequence as this would adversely affect subsequent experiments with intact HBV genomes **(Fig.3A)**. Mutation of either CTCF BS1 (BS1m) or BS2 (BS2m) in isolation or in combination (BS1/2m) abrogated the increase in EnhI activity following CTCF depletion **(Fig.3E)**. Together, these data suggest that CTCF binds to both motifs within EnhI to repress its activity.

**Figure 3:**
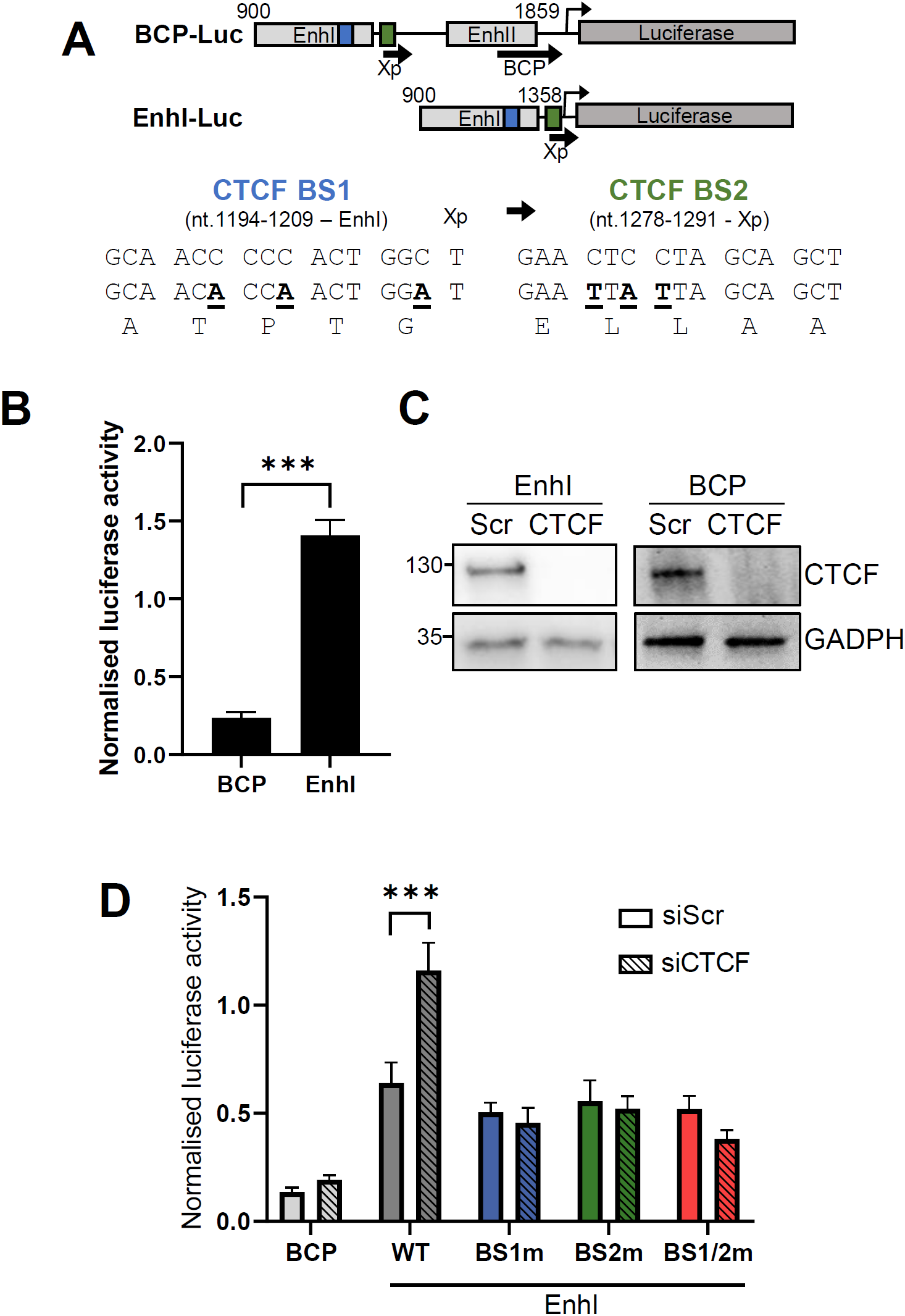
CTCF represses HBV Enhancer I activity. (A) Depiction of HBV genome regions cloned upstream of Firefly luciferase in transcriptional reporter plasmids and mutagenesis strategy of CTCF BS1 and BS2 showing viral enhancers, Xp and BCP, and CTCF BS 1 (blue) and CTCF BS 2 (green). (B) Activity of pEnhI-Luc and pBCP-Luc reporters in HepG2-NTCP cells normalized to co-transfected *Renilla* luciferase expression plasmid. (C) Western blot showing depletion of CTCF following siRNA transfection in pEnhI-Luc and pBCP-Luc transfected HepG2-NTCP cells. (D) Firefly luciferase activity normalized to *Renilla* Luciferase expression in HepG2-NTCP cells co-transfected with pGL3-basic, pEnhI-Luc or pBCP-Luc and either scrambled (Scr) or CTCF-specific siRNA duplexes. (E) Normalized luciferase activity in HepG2-NTCP cells transfected pEnhI-Luc containing mutations in CTCF binding site 1 (BS1m) or 2 (BS2m) or a combination of both (BS1/2m). Data shown are the mean +/- SEM of three independent repetitions. P values were determined by the Sidak’s ANOVA multiple comparisons test. ***denotes p <0.001.

### Silencing CTCF increases HBV preC/pgRNA levels

To determine the effect of CTCF depletion on viral transcripts we selected to use the HepG2-HBV-Epi cells as we previously demonstrated CTCF binding to the viral genome. We confirmed effective knock-down of CTCF at the protein and RNA level 72h post-siRNA transfection **(Fig.4A and B).** We measured HBV RNAs by RT-qPCR as previously described (46) and observed a significant increase in preC/pgRNA levels following CTCF depletion **(Fig.4C)** and an overall increase in total HBV transcripts following CTCF depletion **(Fig.4D).** To determine whether the observed increase in preC/pgRNA levels was due to an alteration of the HBV epigenome following CTCF depletion we measured H4Ac modification of viral DNA as this was previously reported to associate with HBV transcription (47). Silencing of CTCF in HepG2-HBV-Epi cells increased H4Ac abundance within the viral enhancers, BCP Xp and BCP, suggesting that CTCF regulates the epigenetic status of HBV cccDNA **(Fig.4E).**

**Figure 4:**
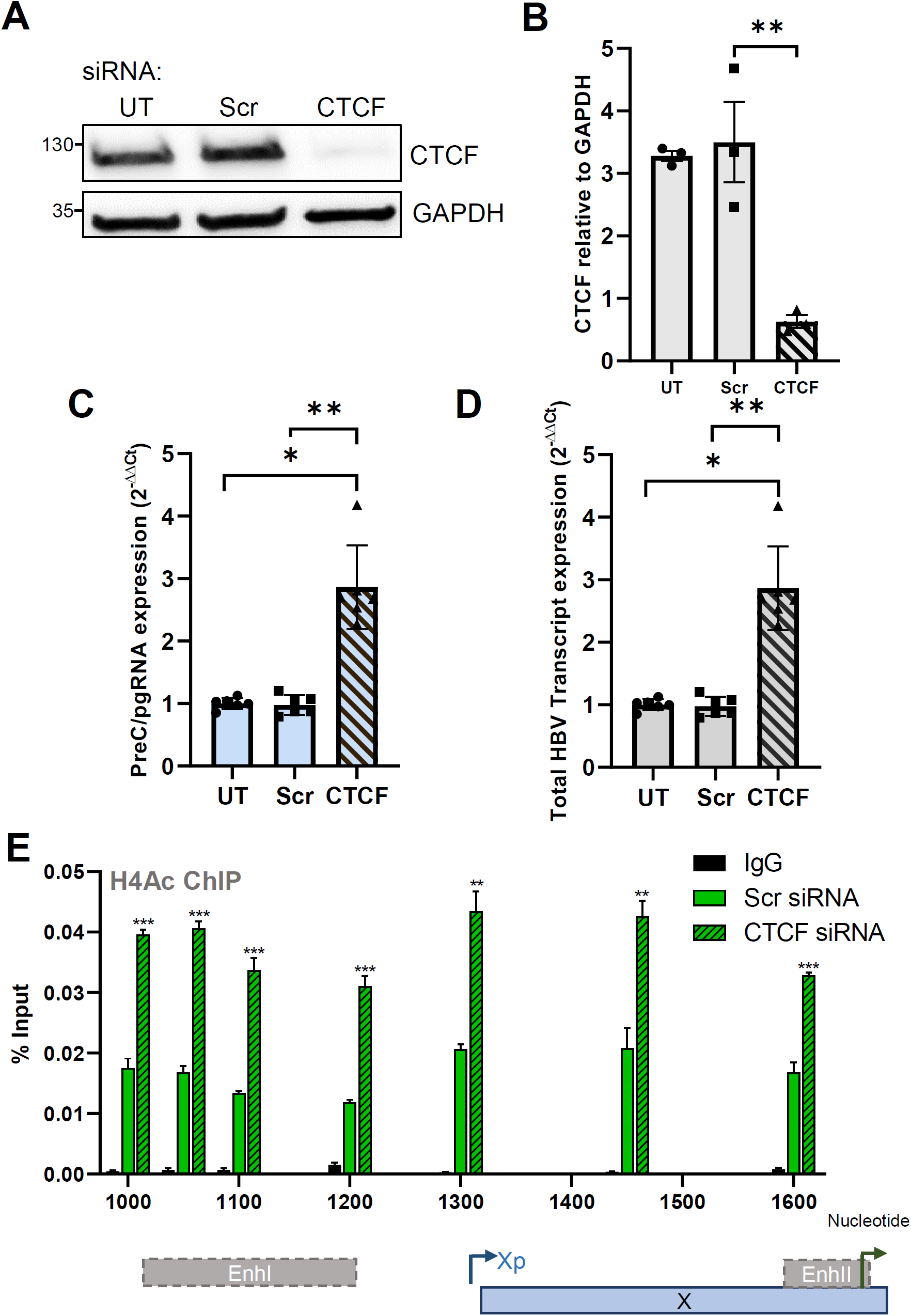
CTCF represses preC/pgRNA transcription from HBV cccDNA. HepG2-HBV-Epi cells were untransfected (UT) or transfected with scrambled (Scr) or CTCF-specific siRNA duplexes and incubated for 72 h. (A) CTCF depletion was assessed by western blotting and quantification in three independent experiments shown (B). (C, D) preC/pgRNA and total HBV RNA abundance were analysed by 4T-qPCR as previously described (46). Data are the mean +/- SD of two independent experiments performed in triplicate. Data are the mean +/- SEM of two independent experiments performed in triplicate. P values were determined by the Kruskal–Wallis ANOVA multiple group comparison. (E) Enrichment of H4Ac marks was assessed by ChIP-qPCR and shown as % Input recovery. P values were determined using a paired t test. *denotes p <0.05, **denotes p <0.01, ***denotes p <0.001.

Hepatocytes are non-proliferating in the healthy liver and most reports studying HBV infection *in vitro* use dimethyl sulfoxide (DMSO) to arrest cells (48). As DMSO has pleiotropic effects on host gene expression (49, 50) we were interested to assess the effects of DMSO on CTCF expression. We noted a significant reduction in CTCF protein levels in DMSO treated cells **(Sup Fig.S1).** We therefore studied the role of CTCF in HBV transcription in non-DMSO treated HepG2-NTCP cells where our protein of interest is abundant.

To extend our studies and to validate a role for CTCF to repress viral transcription during a *de novo* infection, we silenced CTCF in HBV infected HepG2-NTCP cells **(Fig.5A)**. Efficient depletion of CTCF was demonstrated by western blotting **(Fig.5B)** and viral RNAs were analysed by RT-qPCR. In agreement with our earlier data with HepG2-HBV-Epi cells, CTCF depletion in this *de novo* infection model increased preC/pgRNA levels and total HBV RNA **(Fig.5C and D).** Moreover, no significant differences were observed in preS1, preS2 and HBx RNAs **(Fig.5E**). These data support a model where CTCF represses HBV cccDNA transcription, the major transcriptional template in *de novo* infected HepG2-NTCP cells. Taken together, our findings provide evidence that CTCF represses the BCP activity and hence preC/pgRNA levels.

**Figure 5:**
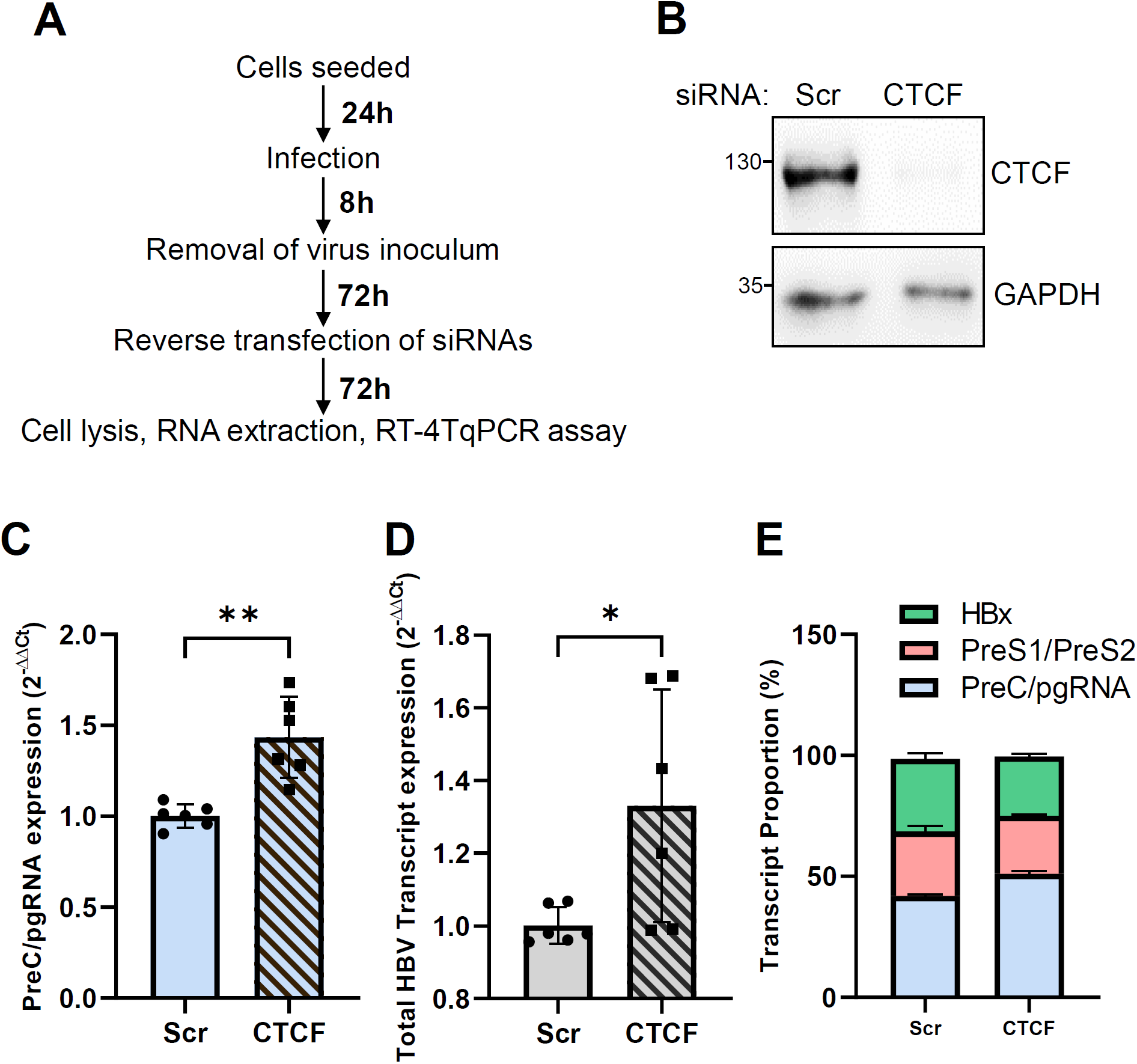
CTCF represses HBV preC/pgRNA transcription in *de novo* infected HepG2-NTCP cells. (A) HBV infected HepG2-NTCP were transfected with scrambled (Scr) or CTCF-specific siRNA duplexes and cultured for 72 h. (B) CTCF depletion was assessed by western blotting and (C) viral transcript abundance analysed by q4T-PCR as previously described (46). Data are the mean +/- SD of two independent experiments performed in triplicate. P values were determined using the Mann-Whitney test (two group comparisons). *denotes p < 0.05, **denotes p < 0.01.

### Mutation of CTCF binding sites within HBV Enhancer I increases transcription

To demonstrate a direct role for CTCF binding to and regulating cccDNA transcription we utilised the HBV minicircle (mcHBV) technology, that recapitulates HBV cccDNA transcription and replication (51). We mutated CTCF BS1 and BS2 alone or in combination in the mcHBV as described in **Fig.3A**. HepG2-NTCP cells were transfected with wild type mcHBV (WT) or mutant mcHBV; BS1m, BS2m or BS1/2m, and harvested 3 days post transfection **(Fig.6A).** Analysis of CTCF binding by ChIP revealed that mutation of BS1 or BS2 alone significantly reduced CTCF binding by over >75% with the combined mutation resulting in an almost complete loss of CTCF-mcHBV complexes **(Fig.6B)**. qPCR analysis showed a significant increase in preC/pgRNA levels when either or both of the CTCF BS were mutated **(Fig.6C)**. However, no differences were observed in HBV mcDNA levels, confirming comparable transfection efficiencies **(Fig.6D)**. These data provide strong evidence of direct recruitment of CTCF to HBV DNA and show the repressive role for CTCF in regulating HBV transcription.

**Figure 6:**
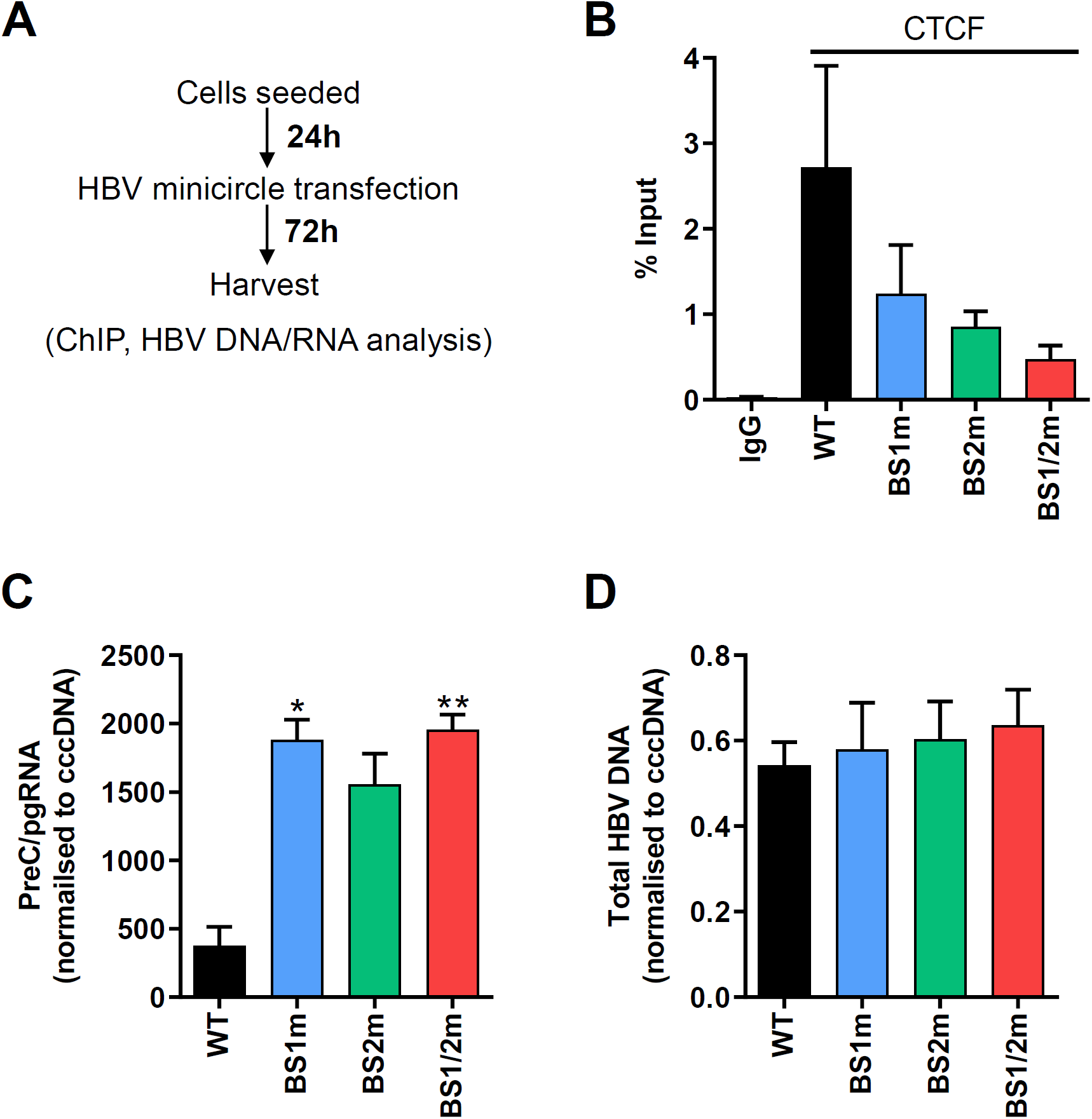
Mutation of CTCF binding sites in HBV mcDNA results in increased preC/pgRNA levels. (A) HepG2-NTCP cells were transfected with wild type HBV mcDNA (WT) or mcDNA with CTCF binding 1 (BS1m) or 2 (BS2m) or both sites mutated in combination (BS1/2m). (B) Cells were harvested 72 h post transfection and CTCF binding analysed by ChIP-qPCR and presented as % of enrichment relative to input chromatin. preC/pgRNA (C) and total HBV DNA (D) levels were quantified by qRT-PCR and normalized to cccDNA amount per cell to account for mcHBV transfection efficiency. Data are the mean +/- SEM of at least three independent experiments. P values were determined using the Kruskal–Wallis ANOVA multiple group comparison. *denotes p < 0.05, **denotes p < 0.01.

## DISCUSSION

In this study we identified two CTCF-binding motifs within transcription regulatory elements, EnhI and Xp, of the HBV genome. We demonstrate CTCF binding to HBV DNA using various model systems that bear both integrated genomes and a cccDNA pool, or cells exclusively expressing episomal copies of viral DNA. Our sonication method sheared cccDNA-like episomes allowing provisional mapping of CTCF binding sites that were confirmed by mutagenesis studies using promoter reporter constructs and mcHBV DNA. Importantly, these CTCF binding sites are conserved amongst all HBV genotypes and across the wider *hepadnaviridae* family, consistent with an evolutionary conserved role in the replication of these viruses. Finally, we show a role for CTCF to repress HBV transcription.

Using several complementary HBV replication models we show that siRNA depletion of CTCF and mutation of CTCF binding sites significantly increase preC/pgRNA levels, consistent with a role for CTCF in repressing viral transcription. To understand the mechanism of CTCF action we used transcriptional reporter assays and found that silencing CTCF significantly increased EnhI activity. Furthermore, mutating the CTCF BS within EnhI attenuated this phenotype, confirming a direct role for CTCF in regulating EnhI. However, analysis of the full BCP, containing both EnhI and EnhII, revealed that the phenotype of CTCF silencing was lost. It is likely that the attenuation of BCP activity following CTCF silencing is explained by the dominant repressive effects of the NRE within EnhII, highlighting the context dependent activity of CTCF in regulating HBV. However, increased activity of the BCP is observed following CTCF silencing in cells containing the full viral episome, which may reflect differential chromatinization and epigenetic modification of the transcriptional reporters as compared to the full viral episome. Alternatively, the transcriptional elements in isolation are no longer subject to regulation by distal elements contained within the intact episome.

To confirm a direct role of CTCF in repressing HBV transcription, we transfected HepG2-NTCP cells with mcHBV mutated in the CTCF BSs. Although the extent to which we could mutate CTCF BS was limited, to maintain the amino acid sequence of the polymerase, we observed a significant reduction of CTCF binding to mcHBV lacking either BS1 or BS2, or both sites mutated in combination. These studies identify the CTCF BSs within the viral genome and confirm CTCF association with HBV DNA. Consistent with the increased preC/pgRNA levels observed in two HBV replication model systems following CTCF depletion, we observed a significant increase in preC/pgRNA when CTCF BS1 was mutated. A similar increase in preC/pgRNA was observed when CTCF BS2 was mutated although this did not reach statistical significance. While the mutation of both BS showed a significant increase in preC/pgRNA abundance, suggesting these sites do not function in a synergistic manner within this model system.

Integration of the HBV genome into the host genome frequently occurs in persistent infection, presumably due to formation of linear double-stranded HBV DNA during aberrant virus replication (52). HBV genome integration is not part of the productive HBV life cycle and the estimated frequency is relatively low (<1 copy per diploid host genome in infected tissues) (53). However, HBV integration can cause host genomic instability leading to tumour progression through tumour suppressor gene inactivation and/or oncogene activation (54). Oncogenic integration events are thought to provide a growth advantage to cells, inducing tumourigenesis (55). HBV integration occurs at random sites, although a preference for integration within regions of open chromatin has been reported (56). It will be interesting to determine whether integration of HBV DNA into the host results in an alteration of local chromatin interactions and host cell gene regulation by the insertion of a virally encoded CTCF binding site(s), as reported for the human retrovirus, HTLV-1 (57). Such genomic rearrangements could have a dramatic effect on host cell gene expression and contribute to HBV-driven carcinogenesis.

Analysis of the epigenetic status of HBV DNA in HepG2.2.15 hepatoma cells revealed a lack of the repressive H3K27Me3 and enrichment of epigenetic marks associated with active transcription in the Xp and BCP regions, downstream of the CTCF binding sites in EnhI/Xp. Similar enrichment of H4Ac was observed in episomal DNA in HepG2-HBV-Epi cells. These findings are consistent with previous reports studying the epigenetic status of HBV cccDNA in various model systems and liver biopsy samples (19, 20). Silencing of CTCF resulted in an increase in H4Ac abundance in HBV cccDNA, which associates with increased HBV preC/pgRNA levels.

Taken together, these findings suggest that CTCF represses HBV transcription by insulating the BCP from the upstream enhancer element, EnhI. EnhI is an important regulator of all HBV promoters and is essential for viral transcription (58, 59). In support of this, HBV-transgenic mice lacking EnhI are defective in virion production (60). The repression of EnhI by CTCF is likely to have a significant impact on the virus life cycle and reduce particle genesis and thereby limit cccDNA pools. Having identified CTCF as a repressor of HBV, we hypothesised that chronic HBV infection may perturb CTCF expression. However, analysing publically available Affymetrix microarray database (61) we found no evidence for HBV infection to perturb intra-hepatic CTCF transcript levels **(Sup Fig.S2)**.

Analysis of the genomic distribution of CTCF BS in the human genome suggests a similar enhancer-blocking activity of CTCF as numerous CTCF binding loci are situated between known transcriptional enhancers and associated promoter elements (62). Such enhancer blocking activity has been extensively characterised at imprinted loci such as the insulin-like growth factor 2 (IGF2)/H19 locus and in development at the β-globin locus (63, 64). CTCF regulates herpes simplex virus differential transcriptional programmes during the lytic and latent phases of the viral life cycle through its enhancer-blocking activity (38). CTCF has been reported to directly repress transcription via recruitment of the Sin3/histone deacetylase (HDAC) compressor complex resulting in reduced histone acetylation (65) that may explain our observations showing increased H4Ac of HBV DNA following CTCF silencing. Our previous work in HPV demonstrated that CTCF represses transcription by stabilising an epigenetically repressed chromatin loop between the viral proximal enhancer and a distal CTCF binding site. However, this repression was not associated with direct binding of CTCF to the HPV enhancer, suggesting that HBV and HPV have evolved fundamentally different mechanisms of CTCF-dependent transcriptional repression.

## METHODS

### Cell lines and antibodies

HepG2.2.15 (41), HepAD38 (42), HepG2-HBV-Epi (43) and HepG2-NTCP cells (48) were maintained in Dulbecco’s Modified Eagles Medium (DMEM, #31966) supplemented with 10% fetal bovine serum (FBS), 2 mM L-glutamine, 1 mM sodium pyruvate, 50 U.mL^−1^ penicillin/streptomycin and non-essential amino acids (all reagents from Invitrogen, UK). All cells were maintained in a 5 % CO_2_ atmosphere at 37°C. HepG2-HBV-Epi cells were kept at low passage to limit HBV DNA integration. The following primary antibodies were used: anti-CTCF (#61311), anti-H3K4Me3 (#39915), anti-H3K27Me3 (#39155) and anti-H4Ac (#39925) were all purchased from Active motif (UK) and anti-GAPDH (SC-32233) was purchased from Santa Cruz.

### ChIP and quantitative PCR

HepG2.215, HepAD38 cells or HepG2-HBV-Epi cells were fixed with 1% formaldehyde (Sigma Aldrich) for 10 min at room temperature before quenching with 125 mM glycine. Cells were washed with ice cold PBS containing EDTA-free protease inhibitors (Roche) and 5 mM sodium butyrate and frozen at −80°C. Pellets were resuspended in ChIP lysis buffer (Active Motif) supplemented with protease inhibitors and incubated on ice for 30 mins. Cells were dounced 30 times using the tight pestle to release nuclei and centrifuged at 2500 xg for 10 mins at 4°C. The supernatant was removed and discarded. Nuclei were resuspended in shearing buffer (Active Motif) pulse sonicated using a Sonics Vibra Cell CV18 sonicator fitted with a micro-probe at 25% amplitude for 15 min on ice using 30 sec on/off cycles. Chromatin samples were cleared by centrifugation and stored at −80°C.

Sonication of HBV cccDNA was evaluated by conventional PCR amplification of increasing amplicon size using a constant sense primer and anti-sense primers described in **Table 1.** Phenol-chloroform extracted DNA from HepG2-HBV-Epi cells before and after sonication was quantified using a NanoDrop ND-1000 spectrophotometer. PCR reactions included 100 ng DNA, MyTaq Red PCR Mix (Bioline, UK) and 200 nM sense/anti-sense primers and amplification following 35 cycles of 95°C, 15 secs; 55°C, 15 secs; 72°C, 30 secs assessed by agarose gel electrophoresis. Products were visualised using SyBr Green Safe dye (Invitrogen).

**Table 1:**
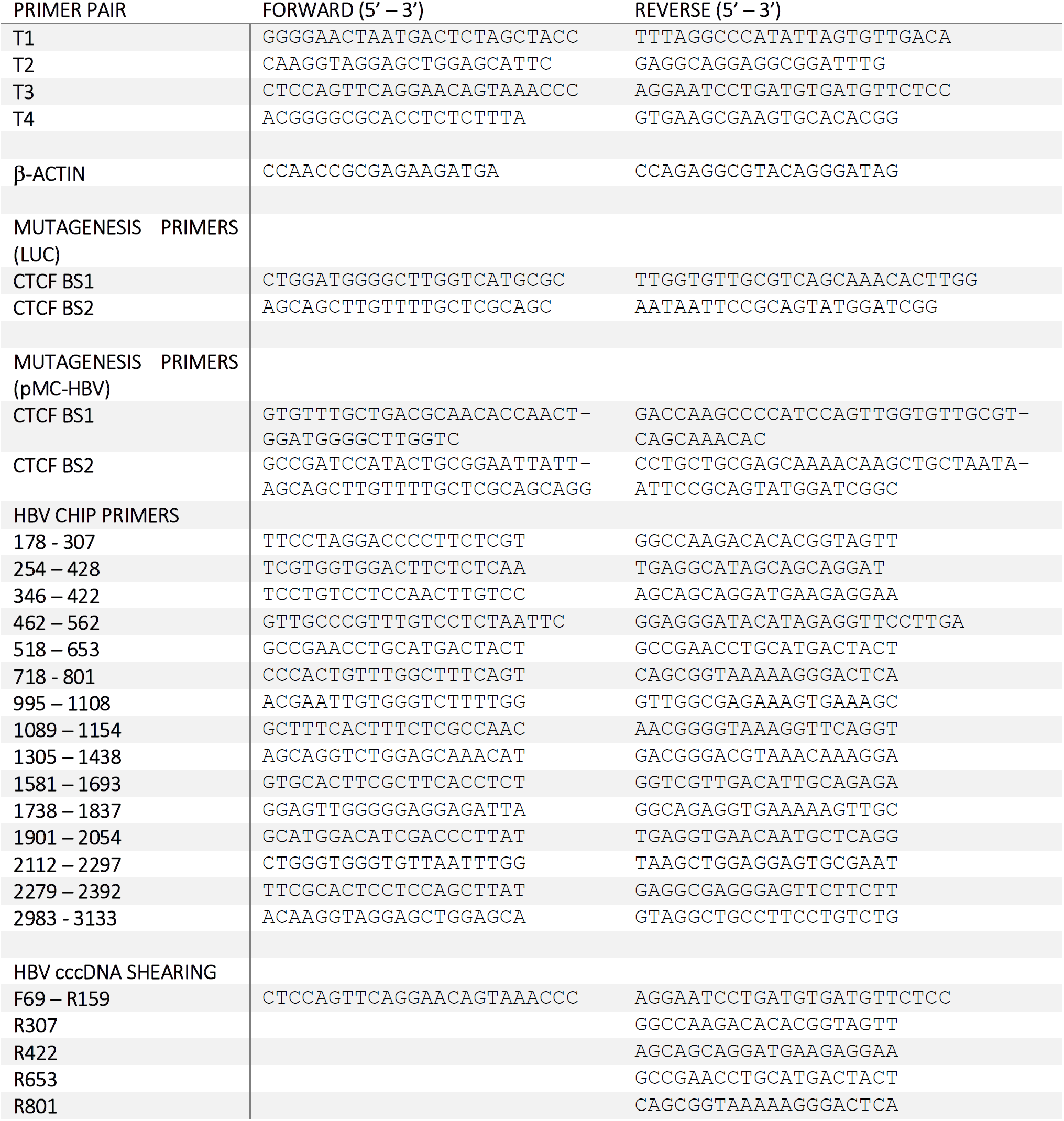
Detailing all primer sequences used.

For ChIP, sonicated lysates were clarified by centrifugation at 16,000 xg for 10 min and CTCF or histone complexes immunoprecipitated with 5-8 µg antibody using a ChIP-IT^®^ Express Chromatin Immunoprecipitation kit, including Protein A magnetic beads as per manufacturer’s instructions (Active Motif, USA). The input and immunoprecipitated DNA were quantified by real-time PCR using a Stratagene MX3500P PCR System. The values were calculated as % recovery respective to input DNA signals. All oligonucleotide sequences are listed in **Table 1.**

### siRNA transfection

Cells were trypsinized to reverse transfect with 25nM of CTCF-specific or scrambled TARGET*plus* Smartpool siRNAs (Horizon, USA) using DharmaFECT4 (20% of amount recommended by the manufacturer’s protocol; Fisher Scientific, Dharmacon). Cells with no siRNA (un-treated; UT) were also assayed to assess lethality of CTCF depletion.

### SDS–PAGE and western blots

Cells were lysed in urea lysis buffer (8M urea, 150mM NaCl, 20 mM Tris, pH 7.5, 0.5 M β-mercaptoethanol) supplemented with protease inhibitor cocktail (Roche) and sonicated for 10 s at 20% amplitude using a Sonics Vibra Cell sonicator fitted with a microprobe. Following quantification of protein concentration by Bradford assay, samples were diluted in Lamelli buffer before incubating at 95°C for 5 min. Proteins were separated on a 10 % polyacrylamide gel and transferred to PVDF membranes (Amersham). The membranes were blocked in TBS-T, 5 % skimmed milk, and proteins detected using specific primary (diluted at 1:1000) and HRP-secondary antibodies (ThermoFisher, diluted at 1:10,000). Protein bands were detected using Pierce SuperSignal West Pico chemiluminescent substrate kit (Pierce) and images collected using a Fusion FX Imaging system (Peqlab).

### HBV transcription reporter assays

1×10^5^ HepG2-NTCP cells were seeded in collagen-coated 24-well plates. Immediately following cell seeding, transfection mixes were added containing 100 ng of either pGL3b-EnhI, pGL3b-BCP or pGL3b-basic, 25 ng *Renilla* luciferase control plasmid (pCMV-Renilla), 25 nM scrambled or CTCF-specific siRNA and 1.5 µl Lipofectamine RNAiMAX™ (ThermoFisher Scientific) in 100 µl OptiMEM (ThermoFisher Scientific). Cells were incubated at 37°C, 5 % CO_2_ for 72 h before being washed with PBS and 200 µL Passive Lysis buffer (Promega, UK) added to each well. Samples were incubated at RT for 30 min with gentle rocking. Lysates were cleared by centrifugation and 20 µL of each added to a white 96-well microtitre plate. FireFly and *Renilla* Luciferase activity were detected using the Dual-Luciferase^®^ Reporter Assay (Promega, UK) using a GloMAX^®^-Multi Detection system (Promega, UK). 50 µL reagent added at a speed of 200 µl/s followed by mixing and 2 s delay. Integration time was 10 s with 1 read/well for Firefly luciferase detection. The same protocol was used for subsequent *Renilla* luciferase detection. Normalised luciferase activity was calculated by dividing Firefly luciferase activity by *Renilla* luciferase activity.

### HBV *de novo* infection

Purified HBV was produced from HepAD38 cells as previously reported (48). HepG2-NTCP cells were seeded on collagen-coated plasticware and infected with HBV at an MOI of 250 genome equivalents per cell in the presence of 4% polyethylene glycol 8,000. Viral inoculum was removed 8 h post infection by extensive washing with PBS and cells maintained in DMSO-free DMEM.

### RNA isolation for cDNA synthesis

Total cellular RNA was extracted using an RNeasy mini kit (Qiagen) following the manufacturer’s protocol. To remove any residual HBV DNA, samples were treated with RNase-Free DNase I (14 Kunitz units/rxn, Qiagen) for 30 min at RT. RNA concentration and quality were assessed using a NanoDrop 1000 spectrophotometer (Thermo Scientific) and 2100 Bioanalyzer (Agilent). cDNA synthesis was performed with 0.25-1 µg of RNA in a 20 µL total reaction volume using a random hexamer/oligo dT strand synthesis kit as per the manufacturer’s instructions (10 min at 25°C; 15 min at 42°C; 15 min at 48°C; SensiFast, Bioline). All oligonucleotide sequences are listed in **Table 1.**

### Quantitative PCR of HBV transcripts

All PCR reactions were performed using a SYBR green real-time PCR protocol (qPCRBIO SyGreen, PCR Biosystems) in a Lightcycler 96™ instrument (Roche). The amplification conditions were: 95°C for 2 min (enzyme activation), followed by 45 cycles of amplification (95°C for 5 s; 60°C for 30 s). A melting curve analysis was performed on the completed reactions to assess specificity and purity of the amplicons (95°C for 10 s; 60°C for 60 s; followed by gradual heating from 60°C to 97°C at 1 °C/s). DNase-treated RNA samples that had not been reverse transcribed were amplified to verify the absence of residual DNA contamination. All oligonucleotide sequences are listed in **Table 1**.

### HBV mcDNA purification and transfection into cells

The plasmid pMC-HBV contains the 1.0 HBV genome (awy) and has been previously described (51). CTCF BS1 and CTCF BS2 were mutated by site-directed PCR mutagenesis using the primers detailed in Table 1 and Prime Star Max (Takara) mutagenesis kit following the manufacturer’s protocols and confirmed by sequencing. ZYCY10P3S2T competent bacteria (System Bioscience) were then transformed with the pMC-HBV (WT, BS1m, BS2m or BS1/2m) and a single colony amplified in Terrific Broth overnight at 37°C. 2 volumes of LB medium supplemented with 0.04 N NaOH and 0.02 % L-Arabinose were added to the culture and further incubated for 8 h at 37°C. Plasmid DNA was extracted using the Nucleobond Xtra Maxi kit according to the manufacturer’s protocol (Macherey-Nagel) and digested with *Nde*I (New England Biolabs) for 2 h at 37°C and plasmid-safe DNase (System Bioscience) overnight at 37°C. After purification, plasmid DNA was assessed by agarose gel electrophoresis to check for elimination of the parental plasmid. HepG2-NTCP cells at 80-90 % confluency were transfected with the pMC-HBV plasmids using TransIT-2020 (Mirus) according to the manufacturer’s protocol in DMEM supplemented with 5 % FBS, 1 % Glutamax and 1 % sodium pyruvate. The following day, cells were washed once with PBS and cultured for 72 h in DMEM supplemented with 5 % FBS, 1 % Glutamax, 1 % sodium pyruvate and 1 % penicillin/streptomycin.

### HBV nucleic acid quantification from mcHBV-transfected cells

Total DNA was extracted using MasterPure™ Complete DNA Purification Kit (Epicentre). Total RNA was extracted using ExtractAll TRI-Reagent (Sigma Aldrich), precipitated in isopropanol, washed in ethanol and resuspended in RNase-free water. Extracted RNA was digested with RNAse-free DNase I (Qiagen) and cDNA synthesised using SuperScript III reverse transcriptase (Invitrogen, Carlsbad, USA). cccDNA was quantified after *Exo*I + *Exo*III endonuclease (Epicentre) digestion of total extracted DNA for 2 hours at 37°C, followed by 20 minutes inactivation at 80°C. Real-time qPCR for total HBV DNA and cccDNA was performed using an Applied QuantStudio 7 machine (BioSystem) and TaqMan Advanced Fast Master Mix. Total HBV DNA was quantified using the TaqMan assay Pa03453406_s1; cccDNA specific primers and probes were: forward 5’-CCGTGTGCACTTCGCTTCA-3’; reverse 5’-GCACAGCTTGGAGGCTTGA-3’ TaqMan probe [6FAM]CATGGAGACCACCGTGAACGCCC[BBQ] (66). Serial dilutions of a plasmid containing an HBV monomer (pHBV-*Eco*RI) served as quantification standard for total HBV DNA and cccDNA. The number of cellular genomes was determined by using the β-globin TaqMan assay Hs00758889_s1 (Thermo Fisher Scientific, Waltham, MA, USA). preC/pgRNA was quantified using the following primers and probe: forward 5’-GGAGTGTGGATTCGCACTCCT-3’; reverse 5’-AGATTGAGATCTTCTGCGAC-3’ and TaqMan probe [6FAM]AGGCAGGTCCCCTAGAAGAAGAACTCC[BBQ] (66). Relative amount was normalized over the expression of housekeeping gene GUSB (Hs99999908_m1, Thermo Fisher Scientific, Waltham, MA, USA).

### Chromatin immunoprecipitation from mcHBV-transfected cells

72h after mcHBV transfection, cells were washed twice with PBS and cross-linked with 1 % formaldehyde for 10 minutes at 37°C. After 5 minutes quenching with 125 mM glycine at 37°C, cells were washed twice with PBS, centrifuged for 5 mins at 300 xg and incubated with Nuclear Lysis Buffer (5 mM PIPES, 85 mM KCl, 0.5% NP-40) for 30 minutes on ice to isolate nuclei. The lysate was then dounced 10 times and centrifuged for 5 minutes at 800 xg at 4°C. Nuclear membranes were then broken by 2 cycles of sonication 30 sec ON, 30 sec OFF on a Bioruptor (Diagenode). Debris were pelleted 10 mins at 11000 xg at 4°C. The supernatant was diluted 10 times with RIPA buffer (10 mM Tris-HCl pH 7.5, 140 mM NaCl, 1 mM EDTA, 0.5 mM EGTA, 1 % Triton X-100, 0.1 % SDS, 0.1 % Na-deoxycholate) supplemented with Complete Mini EDTA-free protease inhibitor (Roche Diagnostics) and 1 mM PMSF and pre-cleared for 2h at 4°C by adding magnetic Protein G Dynabeads (Life Technologies). Beads were discarded and 1 µg of anti-CTCF antibody (Diagenode #C15410210) or isotype matched negative control were added to the chromatin. After an overnight incubation at 4°C, magnetic Protein G Dynabeads and samples incubated for 2 h at 4°C with agitation. Beads were washed 5 times with RIPA buffer, once with TE buffer and resuspended in Elution buffer (20 mM Tris-HCl pH 7.5, 5 mM EDTA, 50 mM NaCl, 1 % SDS, 50 µg/ml proteinase K). Chromatin was reverse crosslinked by incubation at 68°C for 2 h and purified by phenol:chloroform:isoamyl alcohol 25:24:1 (Life Technologies) extraction and ethanol precipitation. cccDNA was quantified using the primers and probes listed above (66).

## Acknowledgements

We acknowledge Stephan Urban (University of Heidelberg, Germany) for providing HepG2-NTCP cells; Ulrika Protzer (TUM, Germany) for providing HepG2-HBV-Epi cells and Wang-Shick Ryu (Yonsei University, South Korea) for HBV promoter-transcription reporter plasmids. We thank Claudia Orbegozo Rubio for expert technical assistance and Chunkyu Ko for advice on HBV biology. BT laboratory is funded by the French Agence nationale de recherche sur le sida et les hépatites virale (ANRS) ECTZ75178. JAM laboratory is funded by Wellcome Trust IA 200838/Z/16/Z and MRC project grant MR/R022011/1. JLP laboratory is funded by MRC project grants MR/R022011/1, MR/T015985/1 and MR/N023498/1. The funders had no role in study design, data collection and interpretation, or the decision to submit the work for publication.

## Author contributions

VDP conducted experiments; JF conducted experiments; GG conducted experiments, FC conducted experiments; PACW conducted experiments; JMH conducted experiments; AC conducted experiments; SB conducted experiments; XZ conducted experiments; PB performed genetic analysis; BT designed experiments and edited manuscript; JAM designed experiments and co-wrote manuscript; JLP designed experiments and co-wrote the manuscript.

## Conflict of interest

None of the authors have any conflict of interest.

## FIGURE LEGENDS

**Supplementary figure 1.**
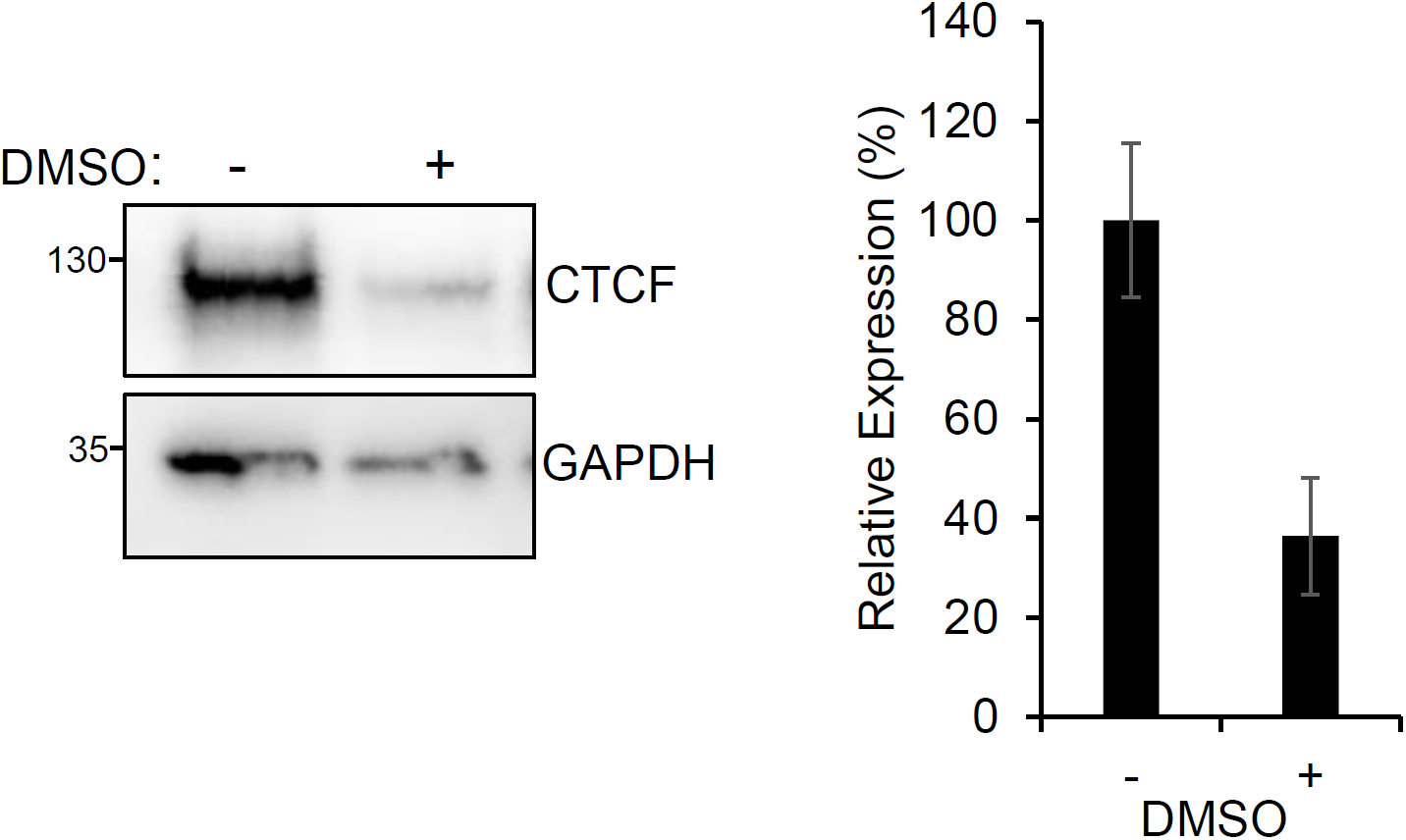
CTCF levels are reduced in DMSO treated HepG2 cells. (A) HepG2-NTCP cells were cultured with (+) or without (-) 2.5 %DMSO for 72 h. CTCF protein levels were assessed by western blotting alongside GAPDH loading control. (B) The relative expression of CTCF compared to GAPDH was quantified by densitometry. Data are the mean +/- SD of three independent experiments.

**Supplementary figure 2:**
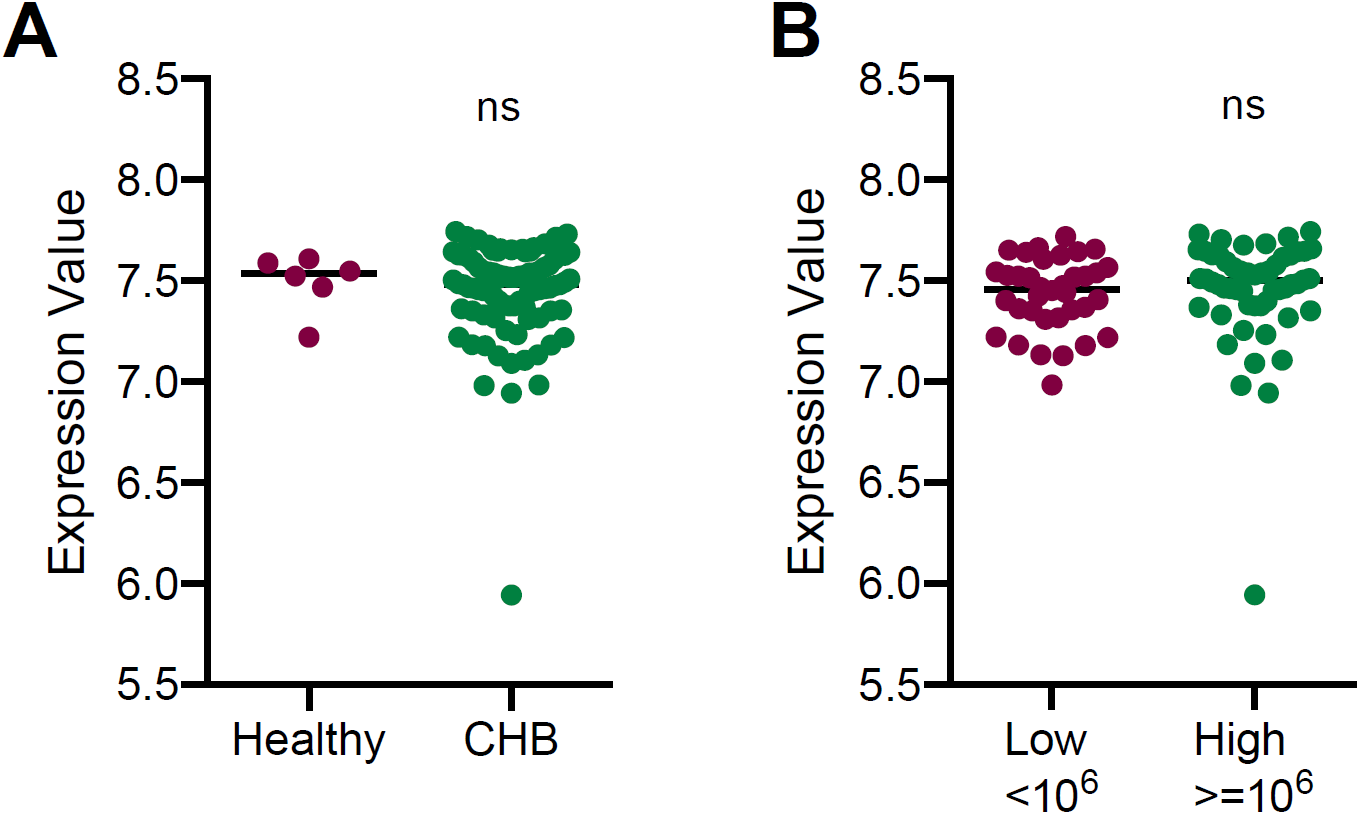
CTCF expression levels in chronic hepatitis B. (A) CTCF RNA levels were determined by high density Affymetrix microarray from liver biopsy samples in non-cirrhotic HBV infected patients (61). Patients with detectable peripheral HBV DNA (n=90) were compared against healthy patient samples (n=6). Statistical analysis was carried out using Mann-Whitney U test. (B) HBV infected patients were categorised into 2 groups based on low (n=36) or high (n=54) peripheral HBV DNA levels, and CTCF expression was compared between the two groups. Statistical analysis was carried out using the Mann-Whitney U test.

